# Preserved CD4^+^ T cell helper function and coordinated antiviral immunity in people with HBV/HIV co-infection on long-term therapy

**DOI:** 10.64898/2026.06.04.730166

**Authors:** Aljawharah Alrubayyi, Anucha Preechanukul, Yinru Zhang, Jonida Kokici, Tanu Kamat, Jessica Davies, Indrajit Ghosh, Fiona Burns, Sabine Kinloch, Pedro Simoes, Sanjay Bhagani, Patrick T F Kennedy, Mala K Maini, Upkar S Gill, Dimitra Peppa

## Abstract

**Background & Aims:** People with HBV/HIV co-infection on antiretroviral therapy achieve higher rates of HBV functional cure than those with HBV mono-infection, yet the immunological basis remains poorly characterised. HBV-specific CD4^+^ T cell responses are critical for viral control and functional cure but have been scarcely examined in HBV/HIV co-infection. Our previous studies in HBV/HIV co-infection demonstrated preserved stem-like CD8^+^ T cells and NK cell functional responses, but whether CD4^+^ T cell helper function is similarly maintained is unknown.

**Methods:** We analysed CD4^+^ T cell responses in 72 participants (HBV n=26, HBV/HIV n=24, HIV n=22) on suppressive antiviral therapy, using multiparameter flow cytometry, virus-specific CD4^+^ T cell functional assays and proliferation assays.

**Results:** People with HBV/HIV co-infection had significantly higher HBV envelope- and core-specific CD4^+^ T cell responses, with IL-2 production particularly discriminating between groups. CD4^+^ T cell responses to CEF (CMV, EBV, and Influenza) were comparable, confirming antigen specificity. Granzyme B-expressing cytotoxic CD4^+^ T cells and TCF-1^+^CD127^+^PD-1^+^CD4^+^ T cells were enriched in co-infection. CD4^+^ and CD8^+^ T cell responses were more frequently coordinated within donors in co-infection than in mono-infection (envelope 83% vs 50%; core 94% vs 60%), where they were more often uncoupled. IL-2 producing CD4^+^ T cells correlated with CD8^+^ T cell responses and the CD4:CD8 ratio in co-infection. HBV-specific proliferative capacity was enhanced in co-infection.

**Conclusions:** People with HBV/HIV co-infection mount functional HBV-specific CD4^+^ T helper responses that are coordinated with CD8^+^ T cell immunity at the individual level. Together with our prior findings of preserved NK and CD8^+^ T cell responses in this cohort, these data identify treated HBV/HIV co-infection as a setting of integrated, rather than compromised, antiviral immunity.

**Impact and Implications:** People with HBV/HIV co-infection can achieve HBV functional cure more frequently than people with HBV mono-infection, but the immune mechanisms remain unclear. This study shows that treated HBV/HIV co-infection is characterised by functional HBV-specific CD4⁺ helper responses and coordinated CD4⁺/CD8⁺ antiviral immunity. These responses were most strongly associated with the CD4:CD8 ratio, a routinely available clinical marker, rather than with CD4 count alone. These findings argue that people with HBV/HIV co-infection should be prioritised in, not excluded from, HBV cure immunotherapy trials.

**Graphical Abstract:** 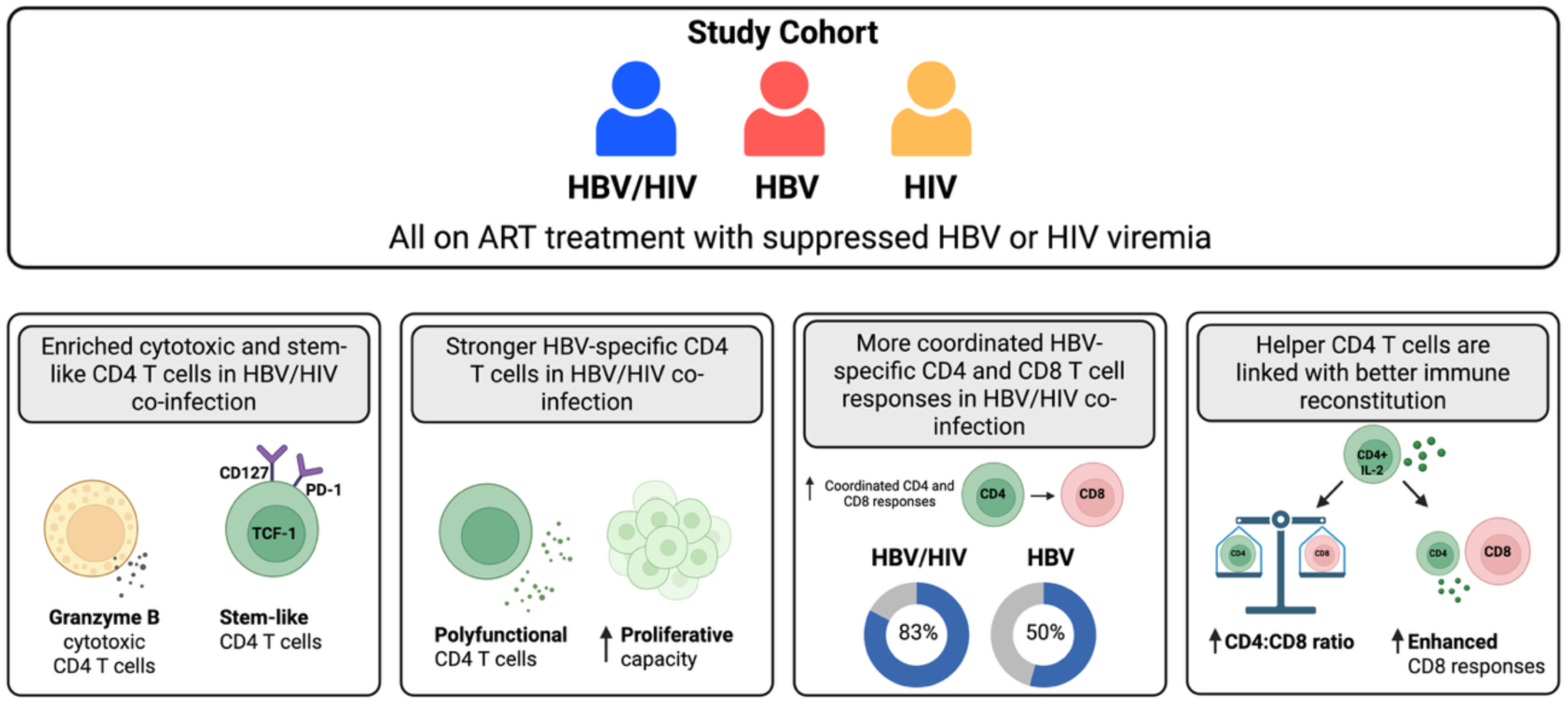

## INTRODUCTION

Approximately 2.7 million of the nearly 40 million people living with HIV are co-infected with hepatitis B virus (HBV).^1^ Despite reported higher rates of liver disease progression in this population,^2^ initiation of tenofovir-containing antiretroviral therapy (ART) is associated with higher rates of HBV functional cure (sustained HBsAg loss) compared with nucleos(t)ide analogue therapy in HBV mono-infection.^3–6^ The immunological mechanisms behind this observation remain poorly understood.^7^ Immune reconstitution following ART and the concurrent reduction in viral antigen burden have been proposed as contributing factors.^6,8^

CD4^+^ T cells are central to HBV control. Animal models have shown that effective HBV clearance requires both CD4^+^ helper and CD8^+^ cytotoxic T cell responses,^9–11^ and recent work demonstrated that CD4^+^ T cells license Kupffer cells to reverse CD8^+^ T cell dysfunction induced by hepatocellular priming.^12^ In a complementary mouse model, CD4⁺ T cell depletion prevented HBsAg seroclearance whereas CD8⁺ T cell depletion did not, and congruent CD4⁺ effector responses were identified in patients achieving clinical cure after nucleos(t)ide analogue withdrawal.^10,13^ In the clinical setting, functional HBV-specific CD4^+^ T cell responses, particularly IL-2 production, distinguish people who achieve a functional cure from those with persistent infection.^11,13^ IL-2 producing CD4^+^ T cells are also predictors of responsiveness to PD-L1 blockade.^14^ Intrahepatic cytotoxic CD4^+^ T lymphocytes (CD4-CTLs) expressing granzyme B, alongside MHC class II-expressing hepatocytes, have been identified as features of functional cure.^15,16^ Together, these findings support that CD4^+^ T cell immunity is a critical determinant of HBV outcome.

Despite this, HBV-specific CD4^+^ T cells remain difficult to characterise. Their frequencies in peripheral blood are extremely low, often at or below the detection limits of standard assays.^17,18^ The CD4^+^ compartment is consequently less studied than CD8^+^ T cells, and data from the HBV/HIV co-infection setting are particularly scarce.^5^ The few existing studies have produced discordant results, likely reflecting differences in assay sensitivity, cohort characteristics, and the era of ART.^19,20^ Much of the existing immunological data from people with HBV/HIV co-infection were generated when lamivudine was the only HBV-active component of ART, tenofovir was not yet available, and treatment was frequently initiated late. More recently, longitudinal data showed that HBsAg decline in people with HBV/HIV co-infection on modern ART regimens was associated with T cell activation and HBV-specific CD4^+^ T cell cytokine production, particularly IL-2 and IFN-γ.^21,22^

We previously reported preserved NK cell antibody-dependent cytotoxicity^23^ and stem-like CD8^+^ T cells with maintained HBV-specific function^24^ in people with HBV/HIV co-infection on long-term tenofovir-based ART. Here, we investigated whether CD4⁺ T cell responses are similarly maintained in HBV/HIV co-infection by assessing global CD4⁺ T cell phenotype, HBV-specific IL-2 production and proliferative capacity, responsiveness to checkpoint blockade, and coordination with CD8⁺ T cell immunity at the individual donor level.

## METHODS

### Study participants

Seventy-two individuals were recruited from the Royal Free Hospital, Mortimer Market Centre, and Barts Liver Centre (London, UK): 26 with chronic HBV mono-infection, 24 with HBV/HIV co-infection, and 22 with HIV mono-infection. All participants were on suppressive antiviral therapy with undetectable HIV and/or HBV viral loads at the time of sampling. People with co-infection were on tenofovir-based ART, while those with HBV mono-infection received either tenofovir or entecavir. Demographic and clinical characteristics are presented in **Supplementary Table 1**. People with co-infection were significantly older (median 56 vs 42 years, p=0.02), had been on antiviral therapy substantially longer (median 15.5 vs 2.1 years, p<0.0001), and had lower circulating HBsAg levels (median 75 vs 274 IU/ml, p=0.009) than those with HBV mono-infection, reflecting the divergent clinical trajectories of the two conditions. ALT levels did not differ between the two groups (p=0.69). People with HBV/HIV co-infection and those with HIV mono-infection were well matched for CD4 count and CD4:CD8 ratio. Sex was recorded as part of demographic data and is reported in **Supplementary Table 1**. The cohort size was not powered for sex-stratified immunological analyses.

This study was approved by the local ethics committee: Berkshire (REC 16/SC/0265), London Bridge (REC 17/LO/0266), and London-Brighton and Sussex (11/LO/0421). Participants gave informed consent to participate in the study before taking part. Peripheral blood mononuclear cells (PBMCs) were isolated by density gradient centrifugation and cryopreserved.

### Phenotypic analysis by flow cytometry

All antibodies used are listed in **Supplementary Table 2**. Cryopreserved PBMCs were thawed and rested in complete RPMI medium (RPMI supplemented with penicillin-streptomycin, l-Glutamine, HEPES, non-essential amino acids, 2-Mercaptoethanol, and 10% FBS). Cells were washed, resuspended in PBS, and surface-stained at 4 °C for 20 min with different combinations of antibodies in the presence of fixable live/dead stain (Invitrogen). Cells were then fixed and permeabilised for the detection of intracellular and intranuclear antigens. The Foxp3 intranuclear staining buffer kit (eBioscience) was used according to the manufacturer’s instructions. Samples were acquired on a BD Fortessa X20 using BD FACSDiva8.0 (BD Biosciences), and subsequent data analysis was performed using FlowJo 10 (TreeStar). Dimensionality reduction was performed on concatenated files using mrc.cytobank platform.

### Analysis of virus-specific CD4^+^ T cell responses and PD-L1 blockade

PBMCs were thawed and rested in complete RPMI. Cells were then stimulated overnight with 3 μg/ml of HBV capsid/core peptides (JPT Peptide Technologies, Catalogue# PM-HBV-CP; 44 peptides, 15-mers overlapping by 11 amino acids), HBV envelope peptides (JPT Peptide Technologies, Catalogue# PM-HBV-LEPULTRA; 121 peptides, 15-mer peptides), HIV-1 Gag peptides (JPT Peptide Technologies, Catalogue# PM-HIV-GAG; 150 peptides, 15-mer peptides), and CEF (Cytomegalovirus (CMV), Epstein-Barr virus (EBV), and Flu (Influenza)) peptides (Miltenyi Biotec, Catalogue# 130-098-426; 32 peptides), or with 0.005% DMSO as a negative control in the presence of αCD28/αCD49d (1 μg/ml) GolgiStop (containing Monensin, 2 μmol/l), GolgiPlug (containing brefeldin A, 10 μg/ml) (BD Biosciences) and anti-CD107α APC/Cy7 antibody (BD Biosciences, Catalogue# 561343, dilution 1 in 50). After stimulation, cells were washed and stained with anti-CCR7 for 30 min at 37 °C and then surface-stained at 4 °C for 20 min with different combinations of surface antibodies in the presence of fixable live/dead stain (Invitrogen Catalogue # L34957, dilution 1 in 300). Cells were then fixed and permeabilised (CytoFix/CytoPerm; BD Biosciences), followed by intracellular cytokine staining. Samples were acquired on a BD Fortessa X20 using BD FACSDiva8.0 (BD Biosciences), and data were analysed using FlowJo 10 (TreeStar). A full list of antibodies used in the ICS assay is listed in **Supplementary Table 2**. Donors were classified as responders if frequencies exceeded 0.02% after background subtraction. For checkpoint blockade experiments, intracellular cytokine staining was performed in the presence or absence of anti-PD-L1 blocking antibody (Invitrogen, 1 μg/ml), which was added during overnight peptide stimulation.

### Proliferation assay

Cryopreserved PBMCs were thawed, washed with 1× PBS and labelled with CellTrace® Violet (CTV, Life Technologies) at a final concentration of 2.5 μM for 20 min at 37 °C. The staining was quenched with FBS, and cells were washed and resuspended in RPMI supplemented with 10% human AB serum (Sigma), 1 mM Penicillin/Streptomycin and 2mM L-Glutamine. CTV-stained cells were then plated at 2.5×10^5^ in 96-well round-bottom plates and stimulated with 1 μg/ml overlapping peptide pools (as described above). Control conditions included media containing 0.005% DMSO (Sigma). Cells were cultured for 7 days at 37 °C. Culture medium was refreshed at day 4 by replacing half the volume with fresh complete medium. Cells were harvested on day 7, stained with viability dye and surface markers as described above and analysed by flow cytometry. Proliferative capacity was assessed by dilution of CTV dye.

### Polyfunctionality and concordance analysis

Boolean gating in FlowJo defined polyfunctional subsets (IFN-γ, IL-2, TNF-α). Pie charts were generated using SPICE v6.1. For concordance, donors with paired CD4^+^ and CD8^+^ data were classified as responders or non-responders independently for each compartment and HBV antigen (Env or Core) (threshold >0.02% IFN-γ^+^TNF-α^+^). Proportions were compared by Fisher’s exact test for dual responders only.

### Statistical analysis

Prism 8 (GraphPad Software) and R Studio were used for statistical analysis. Mann–Whitney U tests were used for comparisons between independent groups, and Wilcoxon matched-pairs signed-rank tests were used for paired comparisons. Correlations were assessed using Spearman’s rank correlation coefficient. Categorical comparisons were assessed using Fisher’s exact test. P values are indicated in the figures. Statistical significance is shown as *p<0.05, **p<0.01, and ***p<0.001. All tests were two-tailed. Polyfunctionality analyses were performed using SPICE version 6.1.

## RESULTS

### Stem-like CD4^+^ T cells are elevated in people with HBV/HIV co-infection

To explore the effects of HBV and the combined effect of HBV/HIV on CD4**^+^** T cell populations, we first analysed the global CD4**^+^** T cell compartment. Consistent with the impact of HIV on CD4**^+^** T cell homeostasis, people with HBV/HIV co-infection exhibited lower frequencies of CD4**^+^** T cells compared to the mono-infection group (**Figure 1A**). No differences were observed in CD4**^+^** T cell activation (measured by HLA-DR and CD38) or CD4**^+^** T cell differentiation between the groups, indicating a preserved CD4**^+^** T cell compartment in HBV/HIV co-infection (**Figure 1B, Supplementary Figure 1A-B**). In contrast, expression of the inhibitory receptor PD-1 differed between groups, with people with HBV/HIV co-infection displaying a significantly higher frequency of PD-1^+^ CD4^+^ T cells compared with HBV mono-infection (**Figure 1C**). Co-expression analysis of PD-1, CD127 and TCF-1 further demonstrated a significantly increased frequency of TCF-1^+^CD127⁺PD-1⁺ CD4**^+^** T cells in the HBV/HIV co-infection compared with HBV mono-infection, suggesting enrichment of a CD4**^+^** T cell population with features of stem-like potential (**Figure 1D**). There were no differences in the global expression of TCF-1 or CD127 between the groups (**Supplementary Figure 1C-D**). Granzyme B^+^ CD4^+^ T cells were enriched in HBV/HIV co-infection and HIV mono-infection compared with HBV mono-infection (**Figure 1E**). These cells are predominantly effector memory CD4**^+^** T cells (**Supplementary Figure 1E**). This is of interest given recent work identifying intrahepatic CD4**^+^** cytotoxic cells as a feature of HBV functional cure.^15,16^ Overall, the coexistence of stem-like and cytotoxic CD4^+^ T cells in co-infection suggests two distinct functional programmes operating within the CD4^+^ T cell compartment.

**Figure 1.**
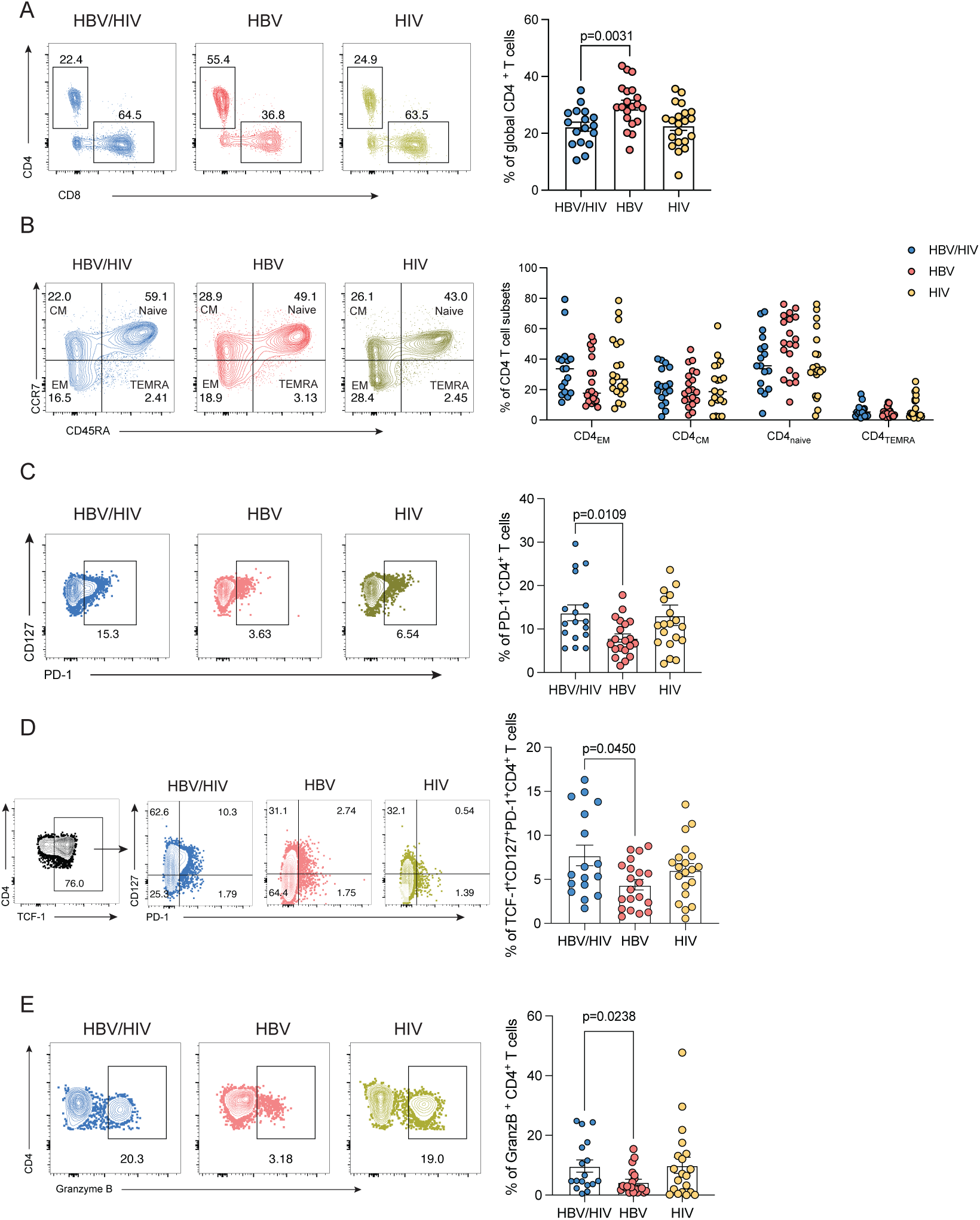
Phenotypic analysis of global CD4⁺ T cells in HBV/HIV co-infection. **(A)** Representative flow cytometry plots and summary data showing the frequency of global CD4⁺ T cells. **(B)** Distribution of CD4⁺ T cell memory subsets defined by CCR7 and CD45RA expression: effector memory, central memory, naïve, and TEMRA. **(C)** Representative plots and summary data showing PD-1⁺ CD4⁺ T cells. **(D)** Representative gating strategy and summary data for TCF-1⁺CD127⁺PD-1⁺ CD4⁺ T cells. **(E)** Representative plots and summary data showing Granzyme B⁺ CD4⁺ T cell frequencies. Mann–Whitney U test was used for comparisons; *p<0.05, **p<0.01.

### Enhanced polyfunctional HBV-specific CD4^+^ T cell responses and proliferative capacity in people with HBV/HIV co-infection

We next assessed whether these phenotypic differences translated into functional responses. After overnight stimulation with overlapping peptides, higher magnitude HBV-specific responses (IFN-γ^+^TNF-α^+^) were observed in HBV/HIV co-infection for both envelope (p=0.0039) and core (p=0.0059) antigens (**Figure 2A-B**). IL-2⁺ CD4⁺ T cell responses were also assessed as a marker of helper function **(Figure 2B).** HBV Env-specific IL-2 responses were significantly increased in people with HBV/HIV co-infection compared with HBV mono-infection (p=0.0087) **(Figure 2B).** These data suggest that HBV/HIV co-infection is associated with enhanced HBV-specific CD4⁺ T cell effector and helper functionality, particularly against HBV-Env. CEF- and HIV Gag-specific CD4⁺ T cell responses were comparable between groups, confirming that the increased responses were specific to HBV antigens **(Figure 2A-B).** HBV-specific CD4^+^ T cells in HBV/HIV co-infection showed greater polyfunctionality, with more cells co-expressing two or more functions (**Figure 2C**).

**Figure 2.**
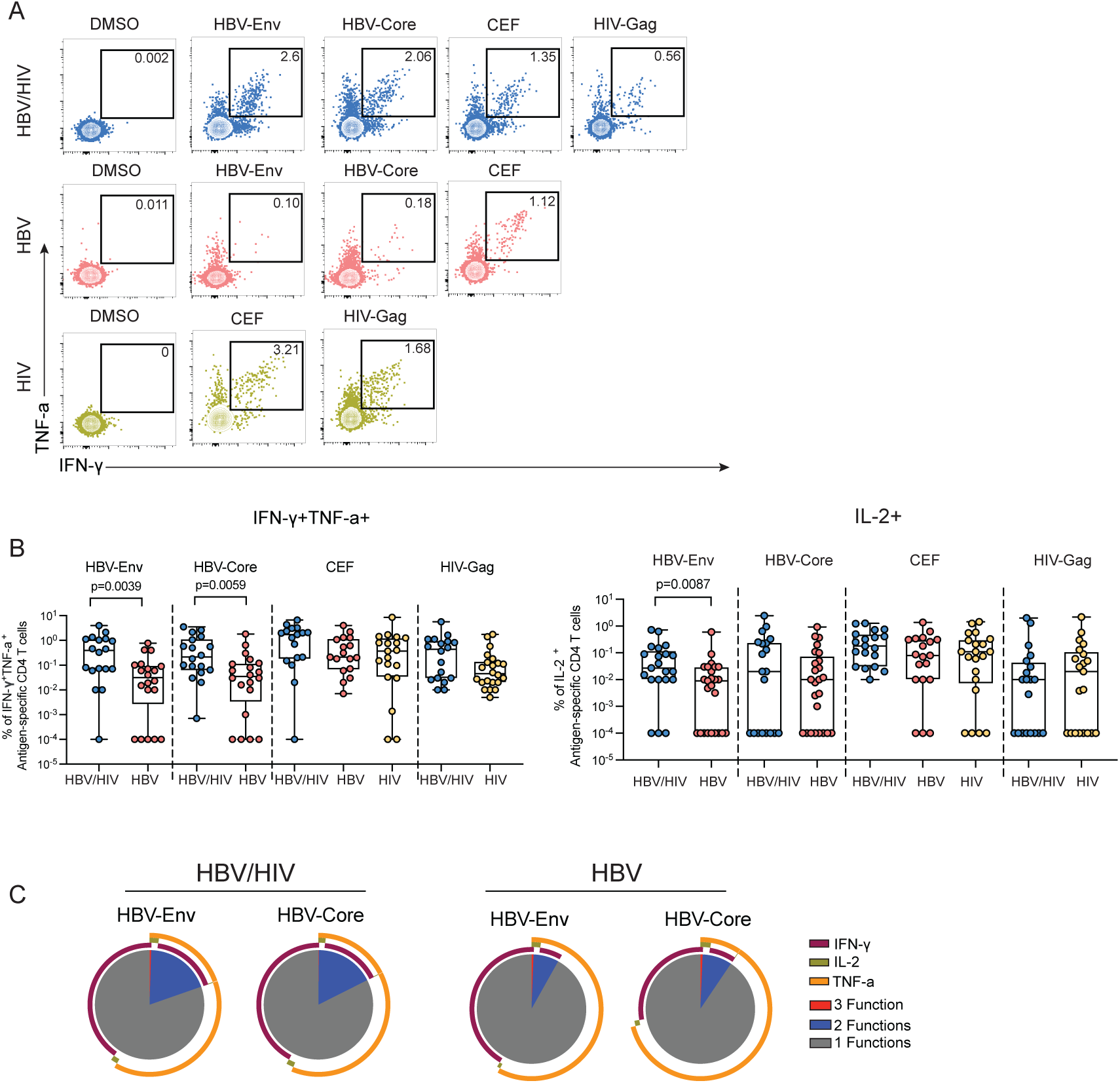
Virus-specific CD4⁺ T-cell responses and polyfunctionality. **(A)** Representative flow cytometry plots showing IFN-γ and TNF-α production by antigen-specific CD4⁺ T cells following stimulation with HBV-Env, HBV-Core, CEF, HIV-Gag or DMSO control. **(B)** Summary data showing IFN-γ⁺TNF-α⁺ and IL-2⁺ antigen-specific CD4⁺ T cell responses across groups. **(C)** Polyfunctional analysis of HBV-specific CD4⁺ T cell responses based on Boolean gating of IFN-γ, TNF-α and IL-2. Mann–Whitney U test was used for pairwise comparisons.

Given the enrichment of stem-like CD4^+^ T cells, we next assessed the proliferative capacity of memory CD4^+^ T cells in people with HBV/HIV co-infection. People with co-infection had significantly higher proliferative responses to HBV envelope and core antigens, as well as CEF peptides (**Supplementary Figure 2A-B**). We also observed a trend toward enhanced HBV-specific CD4^+^ T cell responses after PD-L1 blockade in HBV/HIV co-infection for both Env and Core antigens, whereas responses in mono-infection were unchanged (**Supplementary Figure 2C-D**). This pattern is consistent with our previous findings in CD8^+^ T cells in this cohort ^24^, and suggests that HBV-specific CD4⁺ T cells in HBV/HIV co-infection retain proliferative capacity and checkpoint responsiveness, consistent with a functionally preserved rather than compromised CD4⁺ T cell immunity.

### CD4^+^ and CD8^+^ T cell response concordance is greater in HBV/HIV co-infection

To determine whether CD4^+^ and CD8^+^ T cell responses were coordinated within individuals, we classified donors as dual responders (i.e., having both HBV-specific CD4^+^ and CD8^+^ T cell responses), single responders (having only one response, either CD4^+^ or CD8^+^ T cell response), or non-responders based on IFNγ⁺TNFα⁺ T cell responses for each antigen (Env or Core) independently. Consistent with our previous findings ^24^, HBV-specific CD8^+^ T cell responses were significantly higher in people with HBV/HIV co-infection than in those with HBV mono-infection **(Supplementary Figure 3A**). IL-2-producing CD8^+^ T cell responses were comparable across all study groups **(Supplementary Figure 3A**). Strikingly, for both Env and Core, a larger proportion of people with HBV/HIV co-infection mounted coordinated dual CD4⁺ and CD8⁺ T cell responses than HBV mono-infection (HBV-Env: 83% vs 50% p=0.04; HBV-Core: 94% vs 60% p=0.02; **Figure 3A–B**). In contrast, CEF responses were comparable and high in both groups, indicating that the difference was most evident within HBV-specific immunity rather than reflecting a global defect in T cell responsiveness (94% vs 90%, **Figure 3C**). We next assessed whether HBV-specific CD4⁺ and CD8⁺ T cell response magnitudes were linked within the same individuals. In HBV/HIV co-infection, HBV-specific CD4⁺ and CD8⁺ T cell responses were positively correlated for Env-specific responses (r=0.50, p=0.034, **Figure 3D)** and showed a similar trend for Core-specific responses (r=0.42, p=0.083, **Figure 3F**), whereas no significant correlations were observed in HBV mono-infection for either Env-or Core-specific responses **(Figure 3E, G)**.

**Figure 3.**
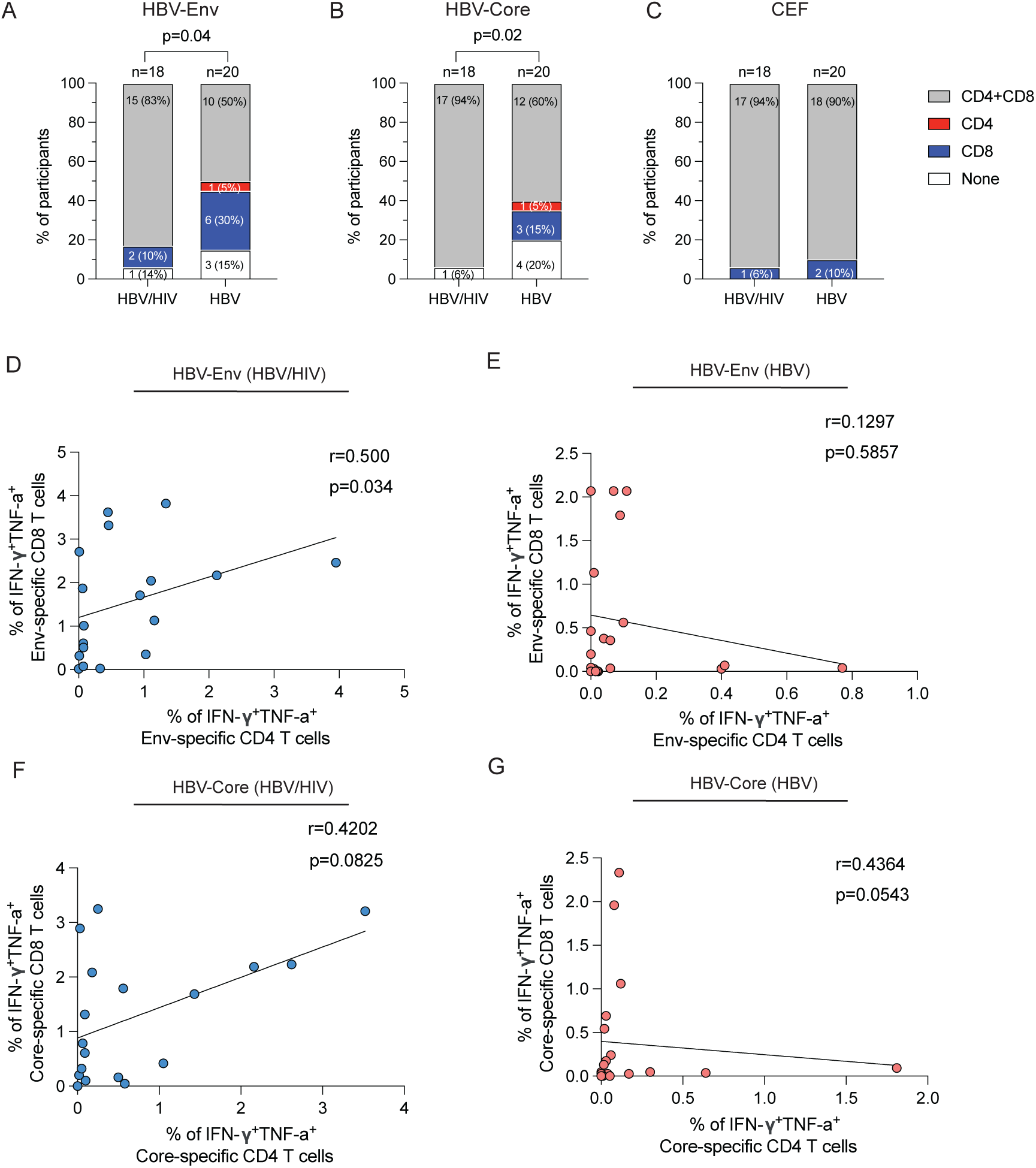
Concordance between HBV-specific CD4⁺ and CD8⁺ T cell responses. (A–C) Stacked bar charts showing the proportion of participants classified as dual responders, CD4 responders, CD8 responders, or non-responders based on IFN-γ⁺TNF-α⁺ responses to HBV-Env **(A)**, HBV-Core **(B)**, and CEF **(C)** in HBV/HIV (n=18) and HBV (n=20). We focused on the participants with data available for both CD4^+^ and CD8^+^ T cell responses to both HBV Env and Core. Fisher’s exact test was used to compare the proportion of dual responders between HBV/HIV co-infection and HBV mono-infection groups. **(D–G)** Spearman correlations between paired CD4⁺ and CD8⁺ T cell responses to HBV-Env in HBV/HIV co-infection **(D)** and HBV mono-infection **(E)**, and to HBV-Core in HBV/HIV co-infection **(F)** and HBV mono-infection **(G)**. Spearman’s rank coefficient was used for correlation analysis.

Having shown that a greater proportion of people with HBV/HIV co-infection mounted dual HBV-specific CD4⁺ and CD8⁺ T-cell responses, we next examined whether this coordination reflected a broader pattern of immune connectivity **(Figure 4A, B**). In HBV/HIV co-infection, stronger HBV-specific CD4⁺ responses correlated with CD8⁺ Env-specific responses within the same donors, supporting a coordinated antiviral T cell response **(Figure 4A)**. We then focused on IL-2-producing CD4⁺ T cells because IL-2 is a key marker of CD4⁺ T cell helper function. Env-specific IL-2⁺ CD4⁺ T cells correlated with Env-specific CD8⁺ T-cell responses in HBV/HIV co-infection (r=0.6314, p=0.0049; **Figure 4C**), but not in HBV mono-infection (r=0.1428, p=0.5597; **Figure 4D**). Core-specific IL-2⁺ CD4⁺ T cells showed a similar positive trend with Core-specific CD8⁺ responses in HBV/HIV co-infection (r=0.4202, p=0.0825; **Figure 4E**), whereas no significant association was seen in HBV mono-infection (r=0.300, p=0.212; **Figure 4F**).

**Figure 4.**
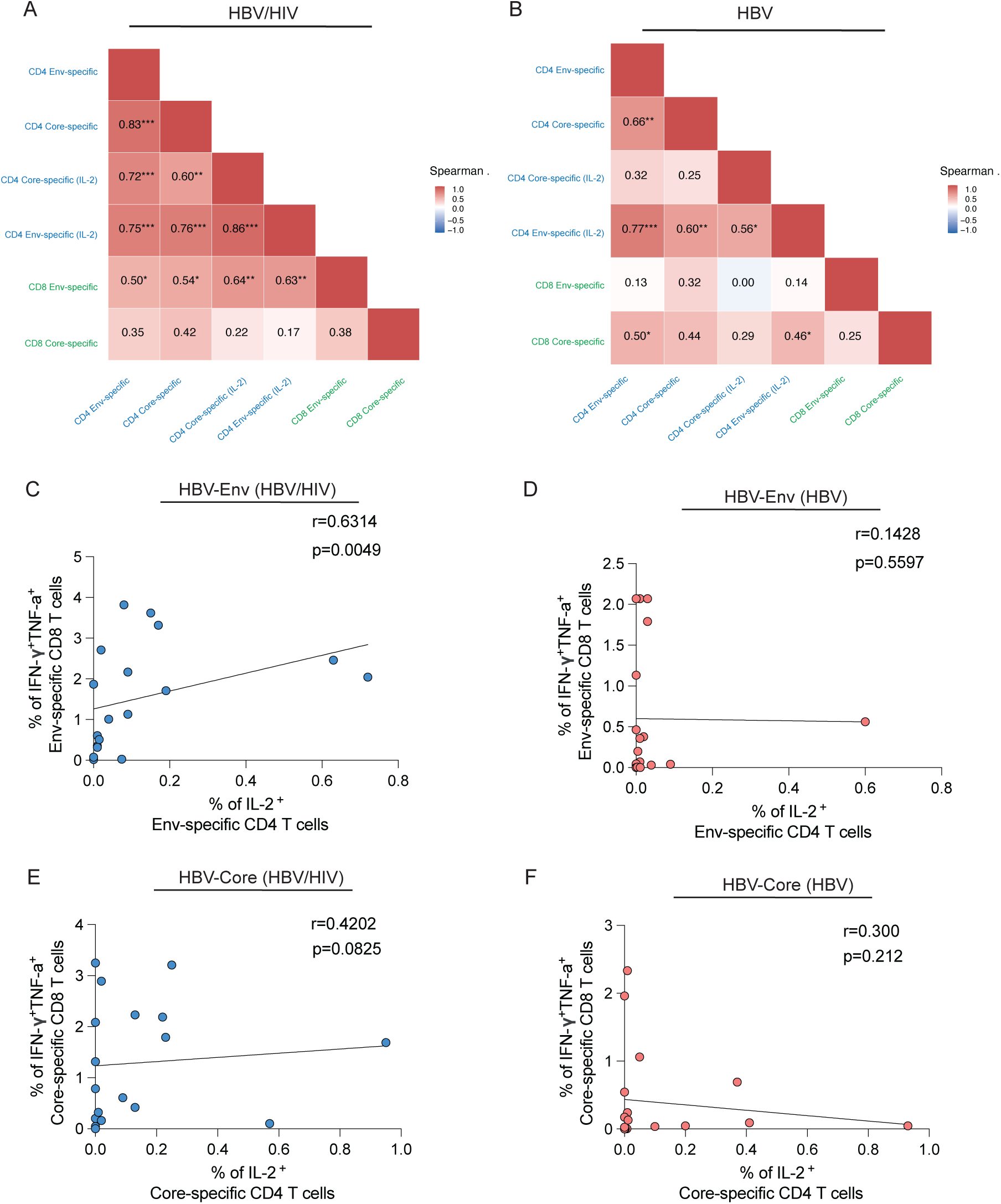
Integrated analysis of HBV-specific CD4⁺ and CD8⁺ T cell function. (A–B) Correlation matrix between HBV-specific CD4⁺ T cell responses and HBV-specific CD8⁺ T cell responses in HBV/HIV co-infection **(A)** and HBV mono-infection **(B). (C–F)** Correlation analysis between IL-2⁺ HBV-specific CD4⁺ T cell responses and IFN-γ⁺TNF-α⁺ HBV-specific CD8⁺ T cell responses for HBV-Env in HBV/HIV co-infection **(C)** and HBV mono-infection **(D)**, and for HBV-Core in HBV/HIV co-infection **(E)** and HBV mono-infection **(F).** Correlations were assessed by Spearman’s rank coefficient. *p<0.05, **p<0.01, ***p<0.001.

### Better immune reconstitution and a higher CD4:CD8 ratio are associated with enhanced HBV-specific CD4^+^ T cell response in people with HBV/HIV co-infection

To explore whether the trajectory of immune reconstitution, rather than a single cross-sectional measure, is associated with HBV-specific immunity, we examined CD4 recovery metrics within the co-infection group. Current CD4 count alone showed no correlation with any functional parameter **(Figure 5A)**. By contrast, the CD4:CD8 ratio showed the strongest association with HBV-specific CD4⁺ T cell responses, correlating with all four functional responses (**Figure 5A**). This association was strongest for IL-2-producing CD4⁺ T cells, both for Core-specific responses (r=0.766, p=0.0002; **Figure 5C**) and Env-specific responses (r=0.738, p=0.0004; **Figure 5E**). Significant correlations were also observed for IFN-γ⁺TNF-α⁺ CD4⁺ T cell responses to Core (r=0.551, p=0.0177; **Figure 5D**) and Env (r=0.534, p=0.0224; **Figure 5F**). Recovery ratio (current CD4 count divided by nadir) showed a more limited association, reaching significance only for envelope IFN-γ^+^TNF-α^+^ responses (r=0.49, p=0.041, **Figure 5A**); CD4 nadir did not correlate with any CD4^+^ T cell functional responses (**Figure 5A**). Treatment duration, HBsAg level, and age did not correlate with any CD4⁺ response parameter within the co-infection group (**Figure 5A**). In mono-infection, none of the available clinical variables correlated with CD4⁺ T cell responses (**Figure 5B**).

**Figure 5.**
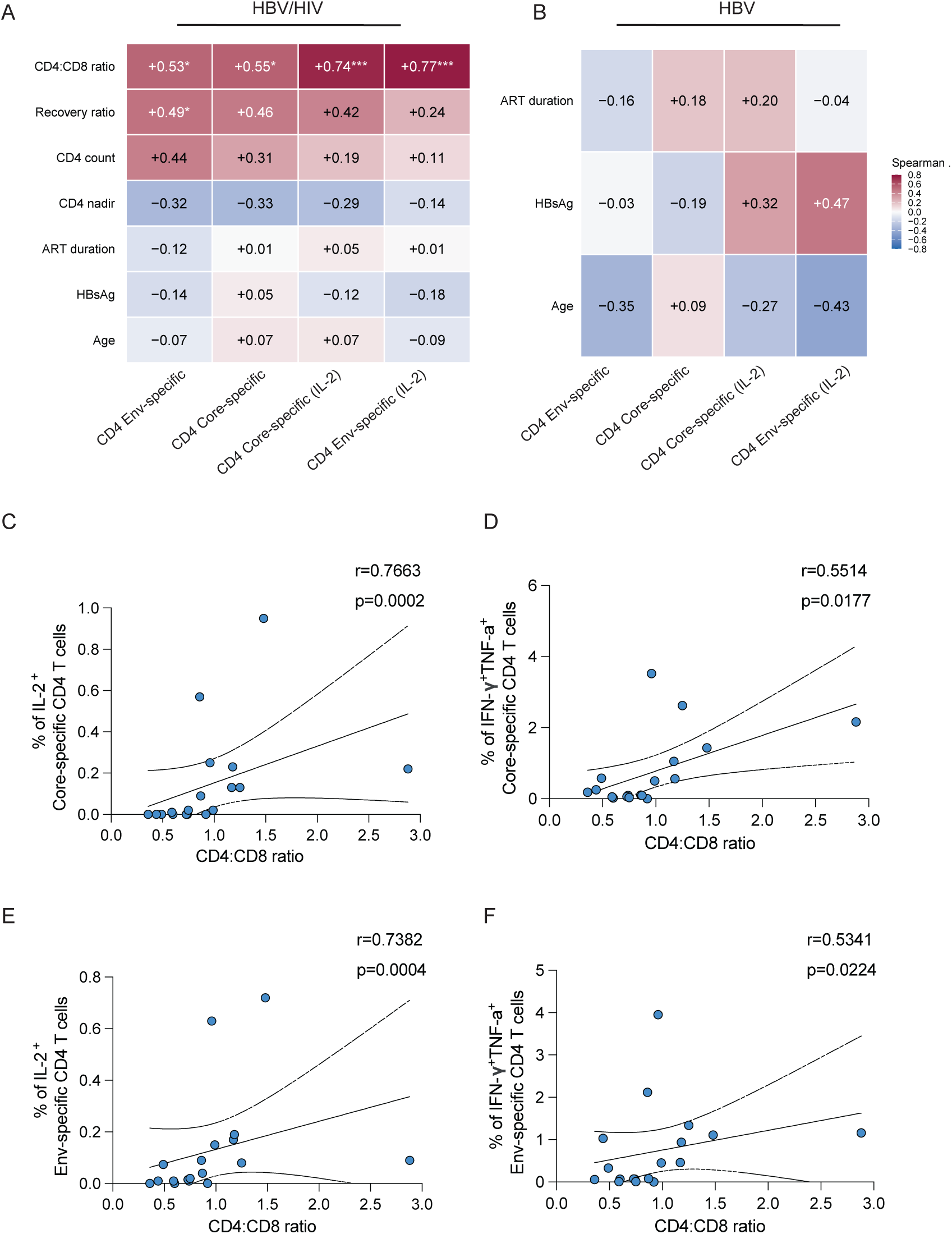
CD4:CD8 ratio is associated with HBV-specific T cell response quality in HBV/HIV co-infection. (A-B) Heatmap showing Spearman correlations between clinical parameters and HBV-specific CD4⁺ functional responses in HBV/HIV co-infection **(A)** and HBV mono-infection **(B)**. **(C-F)** Correlations between CD4:CD8 ratio and Core-specific IL-2⁺ CD4⁺ T cell responses **(C)**, Core-specific IFN-γ⁺TNF-α⁺ CD4⁺ T cell responses **(D)**, Env-specific IL-2⁺ CD4⁺ T cell responses **(E)**, and Env-specific IFN-γ⁺TNF-α⁺ CD4⁺ T cell responses **(F)**. Correlations were assessed by Spearman’s rank coefficient. *p<0.05, **p<0.01, ***p<0.001.

## DISCUSSION

Our findings show that people with treated HBV/HIV co-infection retain a functional CD4^+^ T cell compartment and mount enhanced HBV-specific helper responses with coordinated antiviral responses across both T cell compartments. Together with our prior reports of enhanced NK cell ADCC ^23^ and stem-like CD8^+^ T cell responses^24^ in the same cohort, these findings suggest a pattern of immune preservation in people with well-controlled HBV/HIV co-infection on long-term tenofovir-based ART.

The coordination of CD4^+^ and CD8^+^ T cell responses within individual donors was the most striking feature distinguishing HBV/HIV co-infection from HBV mono-infection. This was not simply a consequence of higher response magnitudes. Virtually all donors in co-infection who mounted a CD4^+^ T cell response also had a concurrent CD8^+^ T cell response, whereas a substantial proportion of donors with mono-infection had CD8^+^ T cell responses without concurrent CD4^+^ T cell help. Recent murine data have shown that stem-like PD-1^+^TCF-1^+^ CD8^+^ T cells are “helpless” cells that require CD4^+^ T cell signals to complete their cytotoxic effector differentiation.^25^ In line with this, CD4⁺ T cell depletion in a mouse model of HBsAg clearance impaired CD8⁺ T cell effector function without reducing CD8⁺ T cell numbers, providing direct evidence that CD4⁺ help augments CD8⁺ T cell effector differentiation in HBV infection.^10^ CD4^+^ help is likewise required for therapeutic HBV vaccine success in preclinical models.^26^ The coordination we observe in co-infection is consistent with this framework, though we cannot establish causality from cross-sectional data.

IL-2 producing CD4^+^ T cells were more discriminating between groups and showed the strongest cross-compartment correlations with CD8^+^ T cell responses. This aligns with the role of IL-2 as a marker of effective HBV-specific CD4^+^ T cell immunity^11,13^ and a predictor of PD-1 blockade responsiveness.^14^ Longitudinal data from a Chinese cohort of people with co-infection on modern ART similarly showed that HBsAg decline was associated with HBV-specific CD4^+^ T cells secreting IL-2 and IFN-γ, and that these associations were stronger for CD4^+^ than CD8^+^ T cell responses.^21^

The CD4:CD8 ratio was the clinical parameter most consistently linked to CD4⁺ T cell responses in HBV/HIV co-infection. The association between CD4:CD8 ratio and response quality was notably stronger for IL-2 producing CD4^+^ T cells, suggesting that the helper arm is most closely linked to the degree of immune reconstitution. These findings extend our CD8⁺ T cell data, where the CD4:CD8 ratio also correlated with HBV-specific responses,^24^ and is consistent with clinical evidence that lower CD4 counts at ART initiation predict higher rates of HBsAg loss.^4,8,27^ CD4 count alone did not correlate with responses, distinguishing this from a trivial cell-number effect. Given its strong association with HBV-specific helper responses and its routine availability in clinical practice, the CD4:CD8 ratio may warrant evaluation as a marker to identify people with HBV/HIV co-infection most likely to respond to HBV cure immunotherapy.

The enrichment of granzyme B-expressing cytotoxic CD4^+^ T cells in HBV/HIV co-infection aligns with recent single-cell analyses of liver tissue from people with functional cure, where intrahepatic CD4-CTLs alongside MHC class II-expressing hepatocytes were identified as features of an altered adaptive immune response associated with HBsAg loss.^15,16^ Cytotoxic CD4⁺ T cells were also expanded in the liver during HBsAg clearance in a recent mouse model.^10^ The coexistence of these cytotoxic effectors alongside IL-2-producing helper cells reinforces the notion that CD4⁺ T cells contribute to HBV control through multiple effector mechanisms.

Several factors may contribute to the distinct immune profile observed in HBV/HIV co-infection. Longer-term HBV suppression and lower HBsAg levels may have reduced chronic antigen stimulation and supported stronger HBV-specific responses^28^, consistent with restoration of HBV-specific T cell function reported after long-term nucleos(t)ide analogue therapy^29^. Better immune reconstitution following long-term ART treatment could further create conditions that support the maintenance or expansion of HBV-specific T cell responses, which may be less evident in HBV mono-infection.^7,8^ This asymmetry is difficult to avoid, as it reflects the divergent clinical trajectories of the two conditions: in Western cohorts, people with HBV/HIV co-infection more commonly acquire HBV in adulthood and initiate HBV-active ART promptly, whereas those with HBV mono-infection often acquire HBV perinatally or in early life, with treatment deferred until disease activity criteria are met.^7,30^ Although treatment duration did not predict T cell phenotype or function within either group, consistent with our previous findings in NK cells ^23^, CD8^+^ T cells, ^24^ and now CD4^+^ T cells, the marked difference in treatment duration remains relevant when interpreting these findings. Notably, the older age of the HBV/HIV co-infection group would be expected to diminish rather than enhance T cell responses, and within-group stratification showed no significant association between HBsAg level and immune parameters. The era of ART must be considered when comparing our findings with earlier studies reporting diminished HBV-specific T cell responses in co-infection.^19,20^ Those cohorts had incomplete HBV suppression, ongoing viral replication, and advanced immunodeficiency. Our cohort comprises people on long-term tenofovir-based ART with sustained dual virological suppression, reflecting contemporary clinical practice. The interpretation of immunological data in HBV/HIV co-infection must account for the complex interplay between infection timing, the stage of chronic hepatitis B at which co-infection occurs, and the treatment context.

### Limitations

This study examines peripheral blood; the extent to which circulating T cell profiles reflect intrahepatic immunity cannot be assumed. The cohort size limited power for within-group stratifications. The cross-sectional design cannot establish causality. Definitive studies would require longitudinal sampling from treatment initiation, which the divergent clinical pathways of the two conditions make logistically challenging. The study was conducted in a Western cohort where infection timing, HBV genotype distribution, and treatment access may differ from settings with higher HBV/HIV co-infection prevalence.

In conclusion, people with virologically suppressed HBV/HIV co-infection have CD4^+^ T cells with functional, coordinated helper responses. The enrichment of IL-2-competent, checkpoint-responsive T cell populations, alongside cytotoxic CD4^+^ T cells, suggests that people with co-infection may benefit from immunotherapeutic interventions targeting HBV cure. These data, together with the companion studies in this cohort, support the inclusion of people with co-infection in HBV cure trials.^31^

## Abbreviations

ADCC: antibody-dependent cellular cytotoxicity
ALT: alanine aminotransferase
ART: antiretroviral therapy
CEF: Cytomegalovirus, Epstein–Barr virus and Influenza
CTL: cytotoxic T lymphocyte
CTV: CellTrace Violet
DMSO: dimethyl sulfoxide
Env: envelope
FBS: fetal bovine serum
HBV: hepatitis B virus
HBsAg: hepatitis B surface antigen
HIV: human immunodeficiency virus
IFN-γ: interferon-gamma
IL-2: interleukin-2
MHC: major histocompatibility complex
NK: natural killer
PBMCs: peripheral blood mononuclear cells
PD-1: programmed cell death protein 1
PD-L1: programmed death-ligand 1
TCF-1: T cell factor 1
TDF: tenofovir disoproxil fumarate
TNF-α: tumour necrosis factor-alpha.

## Acknowledgement

The authors thank all study participants and clinical staff at the Ian Charleson Day Centre at the Royal Free Hospital, Mortimer Market Centre, and Barts Liver Centre.

## Authors’ contributions

AA, performed experiments, analysis and drafting of the manuscript; AP performed experiments and acquisition of data; YZ, JK, and TK, helped with experiments, data acquisition and analysis. JD, IG, FB, SK, PS, SB, PTFK, MKM, and USG contributed clinical samples, data interpretation and critical editing of the manuscript. DP contributed to the conception and design of the study, data interpretation, critical revision of the manuscript and study supervision.

## Ethics approval

This study involves human participants and was approved by the local ethics committee (Berkshire (REC 16/SC/0265) and London Bridge (REC 17/LO/0266)) and conformed to the Declaration of Helsinki principles. Participants gave informed consent to participate in the study before taking part.

## Patient consent for publication

Not applicable.

## Data availability statement

Data are available upon reasonable request to the corresponding author.

