## SUPPLEMENTARY MATERIALS for "Preserved CD4^+^ T cell helper function and coordinated antiviral immunity in people with HBV/HIV co-infection on long-term therapy"

**Supplementary Table 1.** Cohort characteristics for all participants.

**Supplementary Table 2.** List of antibodies used for the phenotypic characterisation.

**Supplementary Figure 1**. CD4$^+$ T cell activation in HBV/HIV co-infection.

**Supplementary Figure 2**. Proliferative HBV-specific CD4$^+$ T cell responses and effect of PD-L1 blockade.

**Supplementary Figure 3**. HBV-specific CD8$^+$ T cell responses in HBV/HIV co-infection.

##### SUPPLEMENTAL FIGURE LEGENDS

**Supplementary Figure 1. CD4$^+$ T cell activation in HBV/HIV co-infection. (A–B)** Representative flow cytometry plots and summary data showing CD38$^+$ CD4$^+$ T cells **(A)** and HLA-DR$^+$ CD4$^+$ T cells **(B)** in people with HBV/HIV, HBV, and HIV infections. **(C–D)** Representative plots and summary data showing global TCF-1$^+$ CD4$^+$ T cells **(C)** and CD127$^+$ CD4$^+$ T cells **(D)**. **(E)** Distribution of memory CD4$^+$ T cell subsets within Granzyme B$^+$ CD4$^+$ T cells.

**Supplementary Figure 2. Proliferative HBV-specific CD4$^+$ T cell responses and effect of PD-L1 blockade.**
**(A)** Representative CTV dilution plots showing antigen-specific CD4$^+$ T cell proliferation following stimulation with HBV-Env, HBV-Core, or CEF in HBV/HIV co-infection and HBV mono-infection. **(B)** Summary data showing proliferative CD4$^+$ T cell responses to HBV-Env,

HBV-Core and CEF across groups. **(C-D)** Representative examples and paired summary data comparing HBV-specific $CD4^+$ T cell responses in the presence or absence of PD-L1 blockade. Mann–Whitney U tests or Wilcoxon signed-rank tests were used as appropriate.

**Supplementary Figure 3. HBV-specific $CD8^+$ T cell responses in HBV/HIV co-infection.** **(A)** Summary data showing IFN-$\gamma^+$TNF-$\alpha^+$ and IL-$2^+$ $CD8^+$ T cell responses following stimulation with HBV-Env, HBV-Core, and CEF in HBV/HIV co-infection and HBV mono-infection. Mann–Whitney U test was used for pairwise comparisons.

**Supplementary Table 1. Cohort characteristics for all participants.**

All patients included are classed as non-viraemic where HBV DNA and/or HIV viral load is below the limit of detection.

| | | **HBV/HIV** | **HBV** | **HIV** | **p value (HBV/HIV) vs (HBV)** |
|---|---|---|---|---|---|
| **Group Size** | n | 24 | 26 | 22 | |
| **Age** | Median (IQR) | 56 (51-60) | 42 (37-56) | 52 (44-58) | 0.0273 |
| **Sex** | Male (n) | 19 | 22 | 18 | 0.721 |
| | Female (n) | 5 | 4 | 4 | |
| **HBV Parameters** | Viral load (DNA IU/ml) | BLD* | BLD* | N/A | N/A |
| | HBeAg status | Positive – 5 Negative - 19 | Positive – 6 Negative – 20 | N/A | 0.848 |
| | ALT, Median (IQR) | 29 (23-37) | 32 (24-40) | N/A | 0.690 |
| | HBsAg (IU/ml), Median (IQR) | 75 (21-297) | 274 (205-882) | N/A | 0.009 |
| **HIV Parameters** | Viral load (RNA copies/ml) | BLD* | N/A | BLD* | |
| | CD4 count, Median (IQR) | 530 (424-661) | N/A | 605 (532-661) | N/A |
| | CD4 nadir, Median (IQR) | 113 (24-234) | N/A | N/A | N/A |
| | CD4:CD8 ratio Median (IQR) | 0.86 (0.59-1.16) | N/A | 0.92 (0.66-1.07) | N/A |
| **Treatment** | Antiviral treatment | Tenofovir based | NUC+ | Tenofovir based | N/A |
| | Treatment duration (years) Median (IQR) | 15.5 (12.25-17) | 2.15 (1.02-6.5) | 6 (3-12) | <0.0001 |

** BLD = Below Limit of Detection during routine blood testing.*

*+ NUC = Nucleos(t)ide analogues (e.g. Tenofovir n = 21, Entecavir n = 5)*

**Supplementary Table 2. List of antibodies used for the phenotypic characterisation.**

| Antibody | Supplier | Cat no. | Clone |
|---|---|---|---|
| BV510 anti-human CD14 | Biolegend | 301842 | M5E2 |
| BV510 anti-human CD19 | Biolegend | 302242 | HIB19 |
| Alexa Fluor 700 anti-human granzyme B | BD Bioscience | 560213 | GB11 |
| BV650 anti-human CD3 | Biolegend | 317324 | OKT3 |
| BV711 anti-human CD8 | Biolegend | 301044 | RPA-T8 |
| PE/Dazzle594 anti-human CD4 | Biolegend | 300548 | RPA-T4 |
| BV785 anti-human CD38 | Biolegend | 303530 | HIT2 |
| BV421 anti-human PD-1 | Biolegend | 329920 | EH12.2H7 |
| Alexa Fluor 700 anti-human CD45RA | Biolegend | 304120 | HI100 |
| FITC anti-human TNF-α | Biolegend | 502906 | Mab11 |
| APC anti-human IFN-γ | Biolegend | 506510 | B27 |
| PerCP eFluor710 anti-human IL-2 | eBioscience | 46702942 | MQ1-17H12 |
| PE-Cy7 anti-human CD154 | Biolegend | 310832 | 24-31 |
| Live/Dead Aqua | ThermoFisher Scientific | L34957 | N/A |
| BB700 anti-human CD4 | BD Bioscience | 566393 | SK3 |
| PE-Cy5 anti-human HLA-DR | BD Bioscience | 562007 | G46-6 |
| PE Texas Red anti-human TIM-3 | Biolegend | 345034 | F38-2E2 |
| PE anti-human LAIR-1 | Abcam | AB269308 | NKTA255 |
| APC-Cy7 anti-human CCR7 | Biolegend | 353212 | G043H7 |
| PE-Cy7 anti-human CD45RA | Biolegend | 304126 | HI100 |
| BV650 anti-human CD127 | Biolegend | 351326 | A019D5 |
| BV605 anti-human CD3 | Biolegend | 317322 | OKT3 |
| FITC anti-human TCF-1 | Cell Signaling Technology | 6444S | C63D9 |
| APC- anti-human TOX | Miltenyi Biotec | 130118474 | REA473 |

### Supplementary Figures:

#### Supplementary Figure 1. CD4$^+$ T cell activation in HBV/HIV co-infection.

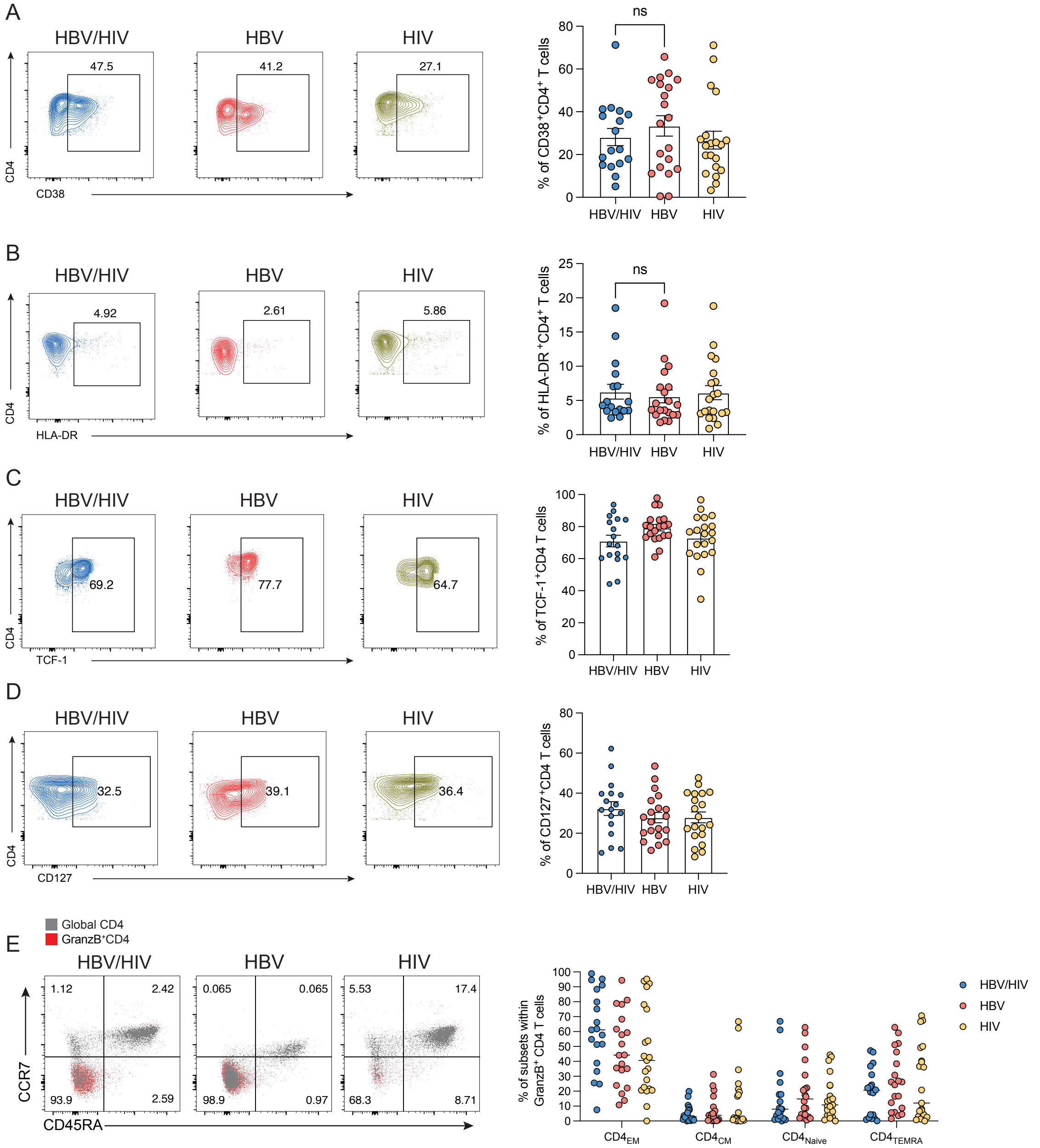

**Supplementary Figure 2. Proliferative HBV-specific CD4⁺ T cell responses and effect of PD-L1 blockade.**

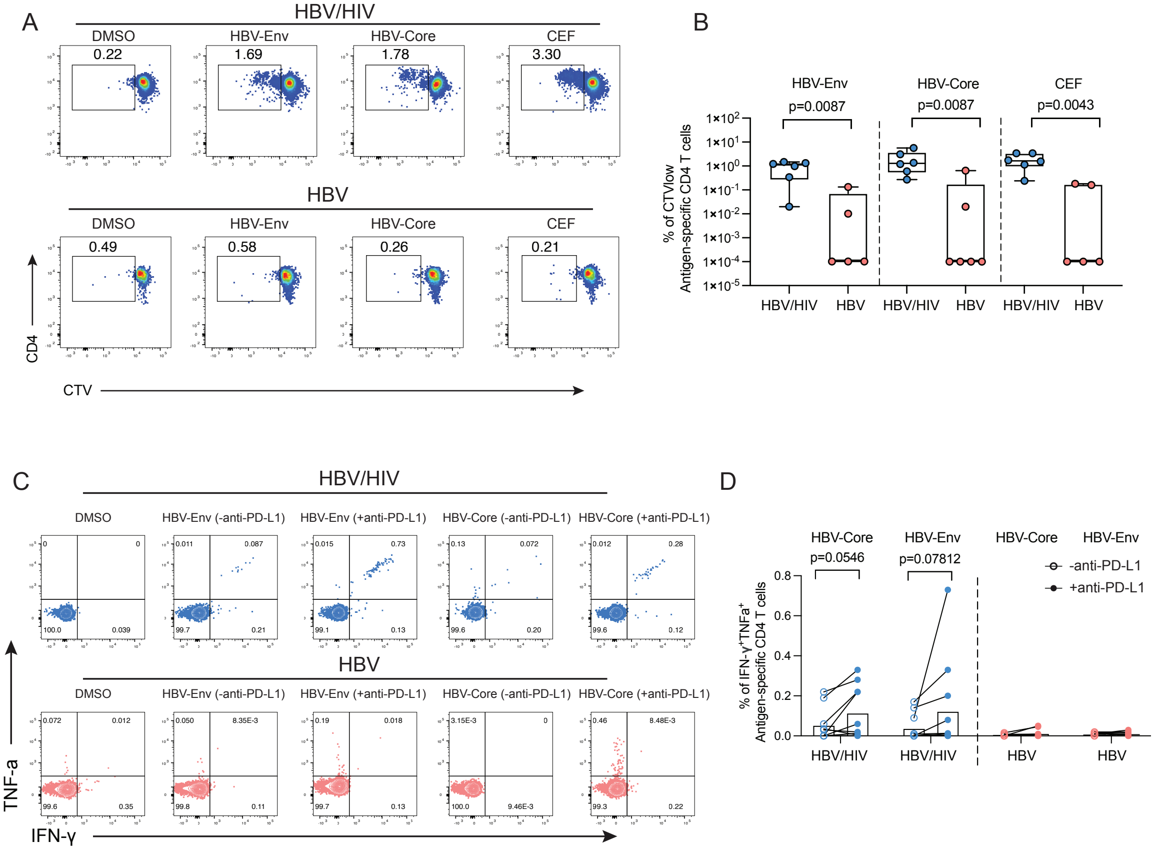

**Supplementary Figure 3. HBV-specific CD8⁺ T cell responses in HBV/HIV co-infection.**

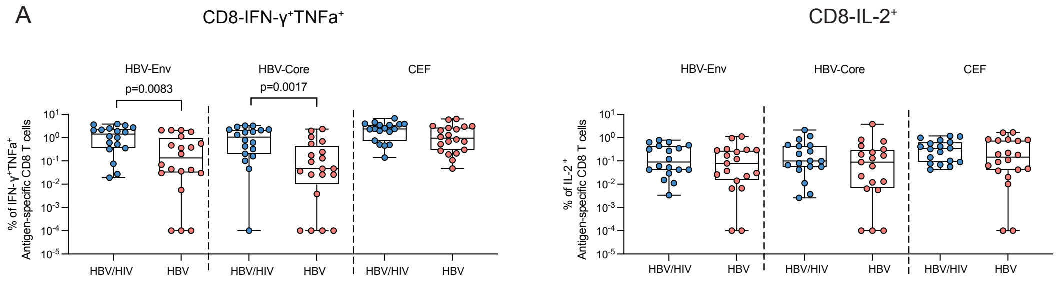
